# A shortcut in forward genetics: concurrent discovery of mutant phenotype and causal mutation in Arabidopsis M2 families via MAD-mapping

**DOI:** 10.1101/2020.06.29.177808

**Authors:** Danalyn R. Holmes, Robert Mobitzer, Markus Wunderlich, Hequan Sun, Farid El Kasmi, Korbinian Schneeberger, Thomas Lahaye

## Abstract

Forward genetics is a powerful tool to establish phenotype-genotype correlations in virtually all areas of plant biology and has been particularly successful in the model plant Arabidopsis. This approach typically starts with a phenotype in an M2 mutant, followed by identifying a causal DNA change in F2 populations resulting from a cross between the mutant and a wildtype individual. Ultimately, two additional generations are needed to pinpoint causal DNA changes upon mutant identification. We postulated that genome-wide allele frequency distributions within the mutants of M2 families facilitate discrimination of causal versus non-causal mutations, essentially eliminating the need for F2 populations. In a proof-of-principle experiment, we aimed to identify signalling components employed by the executor-type resistance (*R*) protein, Bs4C, from pepper (*Capsicum pubescens*). In a native setting, *Bs4C* is transcriptionally activated by and mediates recognition of the transcription activator-like effector AvrBs4 from the bacterial pathogen *Xanthomonas*. Arabidopsis containing an estradiol-inducible *Bs4C* transgene was used in a conditionally lethal screen to identify second-site suppressor mutations. Whole genome sequencing was used for <u>M</u>2 mutant <u>a</u>llele-frequency <u>d</u>istribution (MAD) mapping in three independent M2 families. MAD-mapping uncovered that all three families harboured mutations in *XRN4*, a novel component of executor R protein pathways. Our work demonstrates that causal mutations observed in forward genetic screens can be identified immediately in M2 families instead of derived F2 families. Notably, the timesaving concept of MAD mapping should be applicable to most crop species and will advance the appeal of forward genetics beyond applications in fundamental research.

**SIGNIFICANCE:** Forward genetics has uncovered numerous genes that govern plant immune reactions. This procedure relies on mutant plants with modified immune reactions followed by identification of causal DNA changes in derived F2 progeny. We developed a novel forward genetics concept where causal DNA changes are identified in the initial M2 mutants, making time consuming establishment of F2 populations obsolete. To confirm the feasibility of the concept, we mutagenized transgenic Arabidopsis seeds containing the cell death executing resistance gene *Bs4C* from pepper. Whole-genome sequencing of identified mutant families that lack a *Bs4C*-dependent cell death revealed the *XRN4* gene as a novel component of *Bs4C*-dependent cell death. This confirms our hypothesis that causal mutations can be identified directly within phenotypically selected mutant families.

## INTRODUCTION

To elucidate the molecular basis of biological phenomena, typically the function or activity of key proteins are studied with the intent of uncovering physical or functional connections with other components. Discovery and validation of the components that are involved in a biological process typically involves a synergistic combination of genetic and biochemical approaches. Speed and analytical power are the major parameters when selecting suitable experimental approaches to identify novel elements of a biological process. Forward genetics has been a key discovery tool for biological processes in *Arabidopsis thaliana* (Arabidopsis hereafter) and other plant species, since it requires no prior knowledge of the molecular components that are involved in the process of interest, as it solely relies on differential phenotypes (1). In forward genetics, mutagenesis is often used to induce loss-of-function alleles that typically translate into a phenotypic change in M2 individuals that contain the mutation in a homozygous configuration. Traditionally, causal mutations are located by linkage mapping, usually carried out in F2 populations. Such F2 populations are established by the crossing of M2 individuals to wildtype lines, followed by selfing of the F1. The advent of next generation sequencing (NGS) technologies has drastically simplified this process. Using whole-genome sequencing of bulked DNA of mutant recombinants enabled simultaneous mapping and identification of causal mutations in segregating populations using a single sequencing experiment (2). The base pair resolution of whole-genome sequencing technologies also allowed the use of isogenic crosses (i.e. crosses between the mutant and non-mutagenized individual of the same strain) where random background mutations are used as genetic markers instead of natural DNA polymorphisms between plant genotypes (3, 4). This had the immediate advantage of bypassing practical challenges caused by phenotypic variation between the parental lines of a regular cross that often complicate visual scoring of a specific mutant phenotype in derived segregating populations.

Utilization of isogenic mapping populations also made way for the elimination of two additional generations after the identification of the M2 mutant phenotypes to generate a mapping population. Instead, selfing of heterozygous M2 mutants generates isogenic M3 mapping populations, thereby minimizing the number of generations needed (5). The disadvantage of this method is that it requires the generation of multiple offspring populations since the heterozygous M2 mutants that are needed to establish M3 mapping populations cannot be phenotypically distinguished from wildtype individuals in the M2 generation. As an alternative to genetic mapping, whole-genome sequencing of multiple allelic mutants in one gene outlines a powerful way to identify causal genes without generating any segregating populations (6). While two allelic mutants can already be sufficient for the identification of a candidate gene, this approach is not free from crossing, as the allelism tests relies on pair-wise inter-mutant crosses. Consequently, unless allelic mutants of one gene are known and available, identification of causal DNA changes relies on segregating populations that need to be generated after the identification of the mutant phenotype. This time-consuming and tedious task substantially reduces the appeal of forward genetics. Due to this, only a small fraction of available M2 mutants are usually used for follow up analysis. Therefore, a procedure that does not depend on laborious and time-consuming crosses would enhance the attractiveness of forward genetics.

Plants have two interconnected layers of immunity that collectively provide protection against parasites. Cell surface-localized pattern recognition receptors (PRRs) mediate recognition of conserved pathogen-associated molecular patterns (PAMPs) such as bacterial flagellin (7). To overcome PAMP-triggered immunity (PTI), pathogens have evolved virulence factors known as effectors that are typically translocated into host cells to interfere with PTI and promote disease (8). In response, plants have evolved resistance (*R*) genes that mediate recognition of microbial effectors. Typically, this effector-triggered immunity (ETI) coincides with a plant cell death reaction (hypersensitive response). In most cases, ETI is mediated by intracellular nucleotide-binding/leucine-rich-repeat proteins (NLRs), where they sense activity and/or structural components of microbial effectors and in turn execute a defence reaction (9–11).

Analysis of plant immune reactions triggered by transcription-activator-like effectors (TALEs) from *Xanthomonas* uncovered a mechanistically novel plant *R* gene class where TALEs bind to corresponding effector binding elements within *R* gene promoters and activate transcription of the downstream encoded R protein (12, 13). In such TALE-activated *R* genes, the encoded R protein is not involved in effector recognition, but only in the execution of the plant immune reaction. Accordingly, these R proteins have been designated executors (13, 14).

As of yet, five plant *R* genes that are transcriptionally activated by and mediate recognition of matching TALE proteins have been cloned. With the exception of rice Xa10 and Xa23, which share 50% identity, the predicted executor R proteins show no homology to each other. Recent studies of the executor R protein Bs3 from pepper revealed that the Bs3-triggered immune reaction involves accumulation of salicylic acid (SA), a plant defence hormone that is involved in NLR- and PRR-triggered immune pathways (15). These findings possibly suggest that NLR-, PRR- and executor-type R proteins use, at least in part, common signalling elements to trigger plant defence. However, at this point, little is known about how executor R proteins trigger plant defence.

Bs4C is an executor-type R protein from pepper that was previously shown to mediate recognition of the cognate TALE protein AvrBs4 (16). To identify components of the Bs4C-triggered cell death reaction, we initiated a conditionally lethal forward genetic screen in Arabidopsis that identified three <u>a</u>bolishment of <u>c</u>ell death by <u>e</u>xecutor (*ace*) M2 mutant families. NGS-based <u>M</u>2 mutant <u>a</u>llele-frequency <u>d</u>istribution (MAD) mapping was used instead of commonly used F2 mapping to identify causal mutations and uncovered that all three *ace* mutant families carried mutations in the Arabidopsis *XRN4/EIN5* gene.

## RESULTS

### The pepper executor R protein Bs4C induces plant growth arrest in Arabidopsis

To identify genes that the pepper executor R protein Bs4C requires to trigger plant cell death, we initiated a forward genetic screen in the model system Arabidopsis. To do so, we generated a T-DNA encoding an epitope-tagged Bs4C derivative (*Bs4C-FLAG-GFP*) under the transcriptional control of an estradiol-inducible promoter (Fig. 1*A*) (17). *Agrobacterium tumefaciens* mediated transient transformation of *Nicotiana benthamiana* leaves (agroinfiltration) confirmed that the T-DNA construct mediates cell death in the presence, but not in the absence, of the chemical inducer estradiol, suggesting that the T-DNA construct would confer estradiol dependent *Bs4C* expression in transgenic Arabidopsis plants (Fig. 1*B*). We then transformed the estradiol-inducible *Bs4C-FLAG-GFP* T-DNA (*Estr:Bs4C-FLAG-GFP* hereafter) into the Arabidopsis ecotype Columbia (Col-0 hereafter). We inspected seeds of numerous T2 lines to identify ones that showed a strong, estradiol dependent growth inhibition phenotype. Segregation analysis of T2 seeds on kanamycin containing media identified lines that presumably contain a single-copy transgene insertion. T2 lines with a single-copy transgene and strong seedling growth inhibition phenotype were chosen to produce large quantities of T3 seeds for ethyl methanesulfonate (EMS) mutagenesis. Before carrying out EMS mutagenesis, we confirmed functionality of seedling growth inhibition in T3 seeds. To do so, we placed four-day old seedlings into liquid media containing or lacking estradiol, and analysed seedling growth. We found that in the presence, but not in absence of estradiol, the *Estr:Bs4C-FLAG-GFP* seedlings were severely stunted in their growth (Fig. 1*C*). By contrast, a transgenic line containing a *GFP-GUS* reporter gene under expressional control of the estradiol-inducible promoter (*Estr:GFP-GUS* hereafter) showed no signs of estradiol-dependent growth inhibition. Hence, growth inhibition depends on presence of both the *Bs4C* transgene and estradiol. Immunoblot analysis also showed that the *Estr:Bs4C-FLAG-GFP* transgenic line contained an estradiol-dependent signal matching to the expected 50.6 kDa Bs4C-FLAG-GFP fusion protein (Fig. 1*D*). Taken together, our data illustrate that the pepper executor R protein Bs4C induces cell death when being expressed in the model plant Arabidopsis. Moreover, the established transgenic Arabidopsis lines containing the *Bs4C* gene under control of an estradiol-inducible promoter provide the basis for genetic dissection of Bs4C-dependent cell death in Arabidopsis.

**Figure 1.**
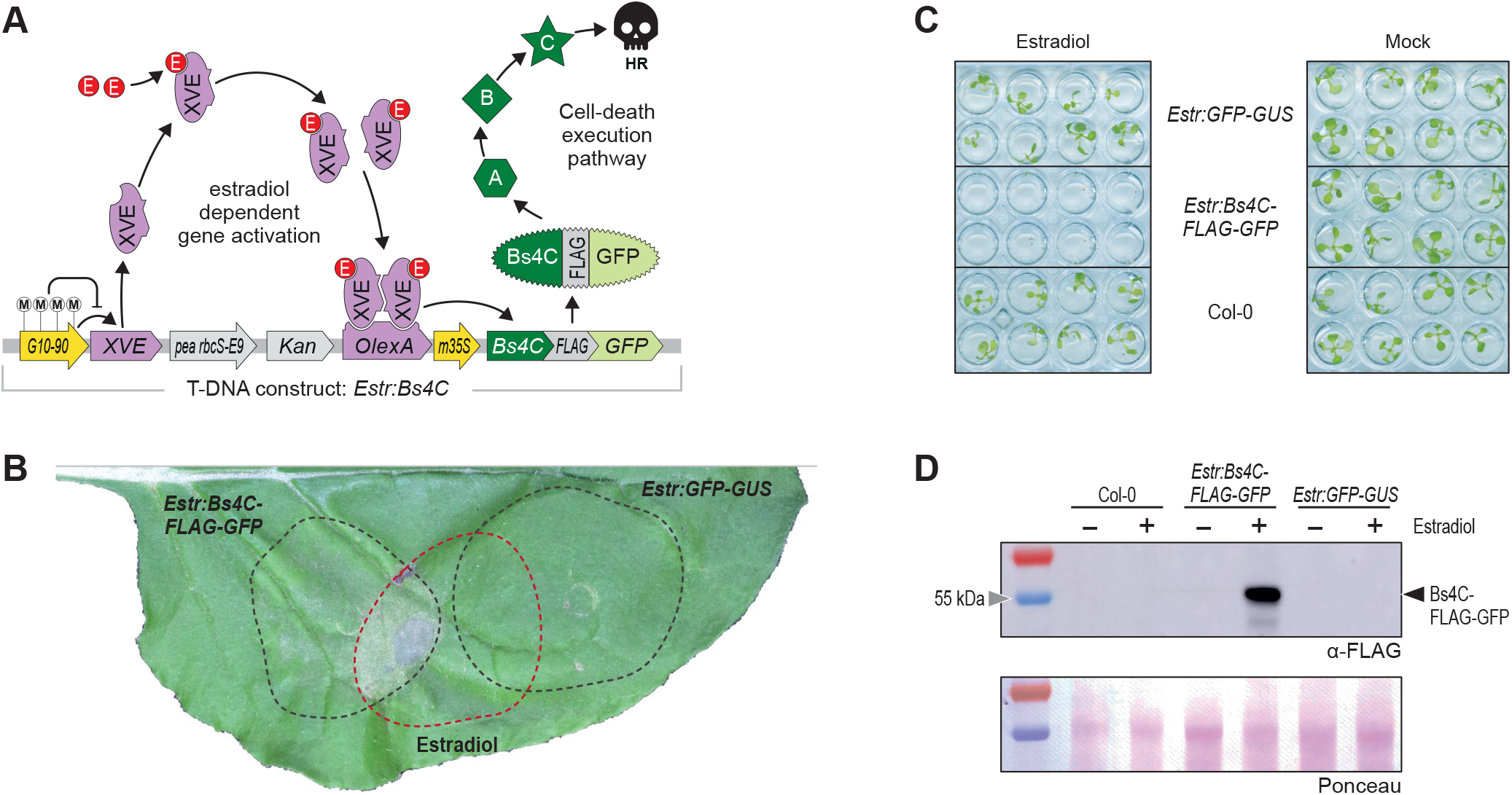
Bs4C induces growth arrest in Arabidopsis. **A** | A T-DNA construct for estradiol-inducible expression of the pepper executor protein Bs4C. Driven by the constitutive G10-90 promoter, the *XVE* gene translates into a chimeric transcriptional activator that contains an estrogen receptor domain. When estrogen (E) is present (here in the form of β-estradiol), it binds to the XVE protein, mediates XVE homodimerization and enables XVE to bind to the LexA operator (OlexA). This induces transcription of the downstream gene encoding a Bs4C-FLAG-GFP protein. The Bs4C fusion protein requires the putative signaling elements A, B, C to trigger plant cell death. Methylation of the G10-90 promoter (M) can cause transcriptional silencing of the G10-90 promoter and results in a non-inducible promoter. **B** | A *Bs4C* transgene triggers estradiol-dependent cell death in *Nicotiana benthamiana leaves*. The depicted T-DNA constructs were delivered into *N. benthamiana* leaves via Agrobacterium mediated transient transformation. Leaf areas into which the inducer estradiol was infiltrated are highlighted with a red line. **C** | An inducible *Bs4C* transgene triggers systemic cell death in Arabidopsis. Four day old seedlings of indicated genotypes were placed in liquid media either containing estradiol or a lacking estradiol (Mock). Ten days later, the seedlings show cell death in presence of estradiol and the *Bs4C* transgene. **D** | Immunoblot analysis using anti-FLAG antibody of tissue from two week old Arabidopsis seedlings of depicted genotypes (Col-0, *Estr:Bs4C-FLAG-GFP*, *Estr:GFP-GUS*). Plants were incubated for 24 hours in liquid media either containing estradiol or lacking estradiol (mock). Ponceau stained membrane serves as a loading control.

### A conditionally lethal screen identifies Arabidopsis mutants that do not execute a Bs4C dependent cell death

To induce randomly distributed mutations across the Arabidopsis genome, approximately 10,000 *Estr:Bs4C-FLAG-GFP* T3 (M0) seeds were treated with EMS and planted into soil. Corresponding M1 plants were individually bagged, and derived M2 seeds were harvested, creating 4,000 M2 families. About 100 seeds of each M2 family, equating to approximately 400,000 M2 seeds in total, were studied as representatives for the entire M2 families. Seeds were allowed to grow on agar plates containing estradiol in an effort to identify second-site suppressor mutants that inhibit Bs4C-dependent cell death. After 14 days, most seedlings had stopped growing and neglected cotyledon emergence (Fig. 2*A*). M2 families containing putative suppressor mutations were easily detectable, as they were large in size and developed roots and true leaves with a green colour similar to *Estr:GFP-GUS* (Fig. 2*B*). A total of 46 M2 families contained individual plants that grew like *Estr:GFP-GUS* plants on estradiol-containing agar plates. A low percentage of survivors within most M2 families suggests recessive inheritance of suppressor alleles. Survivors from each M2 family were transplanted from estradiol plates to soil for further investigation. Here, we present the analysis of three representative *ace* mutant families.

**Figure 2.**
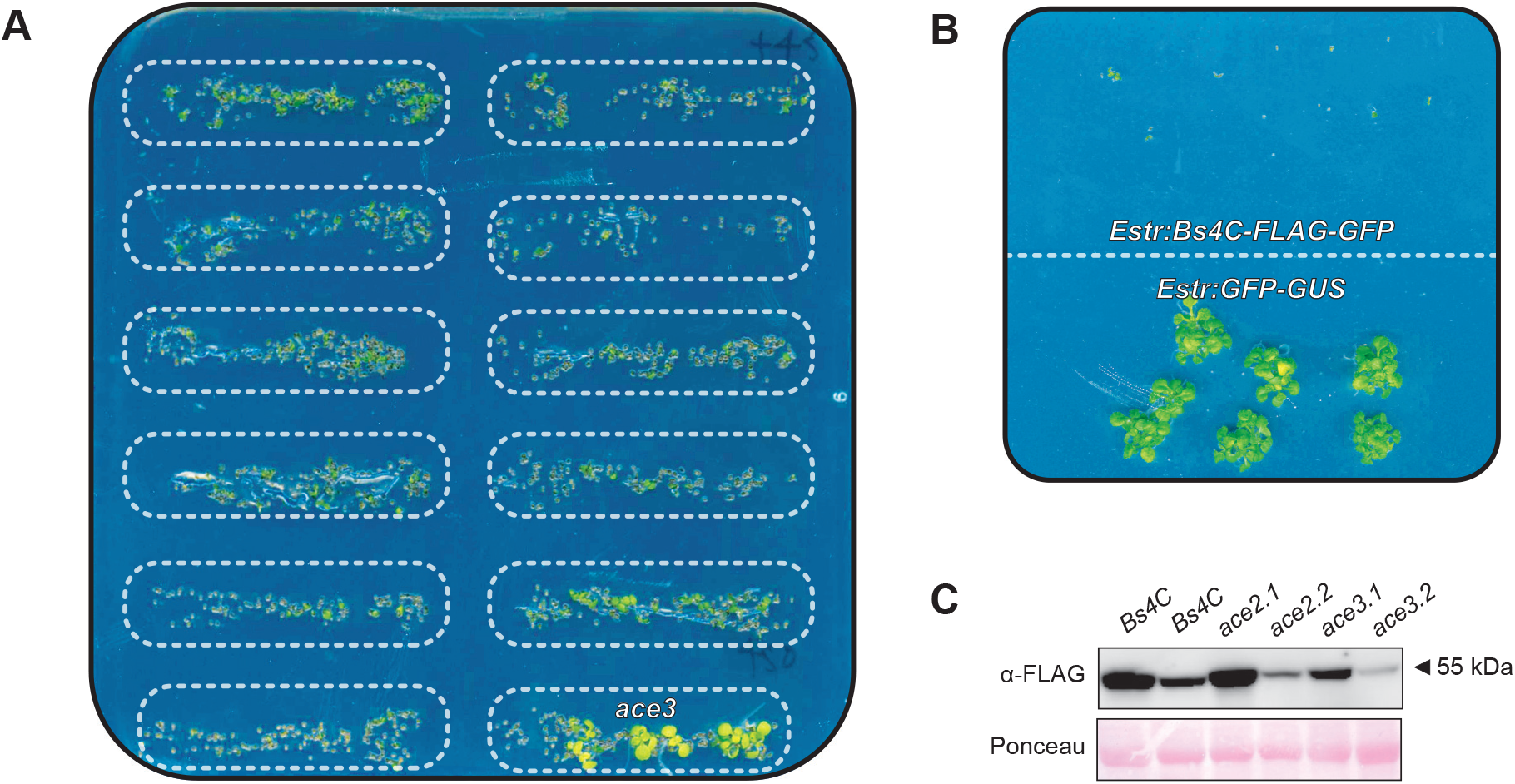
*ace* screen identifies suppressors of Bs4C-dependent cell death in Arabidopsis. **A** | Identification of the *ace3* M2 family. Seeds of twelve distinct M2 families are placed in rows on estradiol-containing agar. Boxes framed by dashed lines indicate the region that is covered by seeds of one M2 family. One M2 family (*ace3*; bottom right) contains individual M2 plants that grow despite the presence of the inducer chemical. **B** | Estradiol triggers a Bs4C-dependent cell death reaction. Seeds containing either an estradiol-inducible Bs4C (*Estr:Bs4C-FLAG-GFP*) or a *GFP-GUS* reporter gene (*Estr:GFP-GUS*) were placed on an estradiol-containing agar plate. **C** | Bs4C protein is expressed in different *ace* mutants. Immunoblot analysis of five week old Arabidopsis leaves treated with estradiol for 24 hours. Bs4C was detected using an anti-FLAG antibody. Ponceau stain provides an info on total protein content in the samples.

### *ace* mutants show no systemic cell death despite having a functional *Bs4C* gene

Two classes of mutations were expected to be identified from our forward genetic screen: those that are within putative signalling and/or regulatory components that Bs4C requires to induce plant cell death, and those that are within the transgene and affect expression and/or functionality of Bs4C. To exclude plants that did not accumulate similar levels of Bs4C protein to that of the parental line, we analysed Bs4C protein expression in *ace1*, *ace2,* and *ace3* mutants by immunoblot analysis. In all three *ace* mutants, immunoblot analysis highlighted signals matching to the expected 50.6 kDa Bs4C-FLAG-GFP fusion protein (Fig. 2*D*, Fig. S1). Moreover, we PCR-amplified and sequenced the *Bs4C* coding sequence (CDS) in all three *ace* mutants’ families and found that they all contained the wildtype *Bs4C* CDS.

### Segregation in *ace* M2 seeds does not fit to the expected 1:7 ratio

Arabidopsis M1 seeds contain two diploid cells that give rise to generative organs (inflorescence) of M1 plants that can be phenotypically studied in the M2 generation (1, 18). If EMS mutagenesis induces a mutation in one of the two diploid M1 precursor cells, this translates into a 1:7 phenotypical segregation of bulked M2 family seeds, assuming recessive inheritance. Thus 12.5% of the seeds of each *ace* M2 family are expected to survive on agar plates containing estradiol. We plated several hundred M2 seeds for each of the three *ace* families on agar plates containing estradiol and observed survival rates of 6.1% (45/734), 2.0% (17/833), and 5.1% (57/1114) for *ace1*, *ace2,* and *ace3* mutant families, respectively. Given the clear phenotype in all three *ace* mutant families and the large number of studied M2 seeds, it seems unlikely that deviations of observed versus expected segregation ratios are due to errors in phenotypical scoring. Thus, the observed distorted segregation is possibly the consequence of diplontic selection, a process of competition between cells within a meristem that can result in reduced proliferation of mutated cells (19).

### Segregation of EMS mutations in *ace* M2 families provides a basis to identify causal mutations

Irrespective of the observed segregation data, such M2 plants that grow in the presence of estradiol (survivors) should have the causal mutation exclusively in the homozygous configuration, assuming recessive inheritance. By contrast, non-causal mutations are expected to segregate randomly in M2 survivors with exception of those that are linked to the causal mutations. NGS-mediated analysis of the allele frequency for each EMS mutation within a pool of survivors from one M2 family can therefore reveal mutations that are homozygous across all M2 survivors in one family and that are potentially causal for the observed cell death suppression phenotype (Fig. 3). We termed this concept as <u>M</u>2 mutant <u>a</u>llele-frequency <u>d</u>istribution (MAD) mapping and studied the feasibility of MAD-mapping in three representative *ace* mutant families.

**Figure 3.**
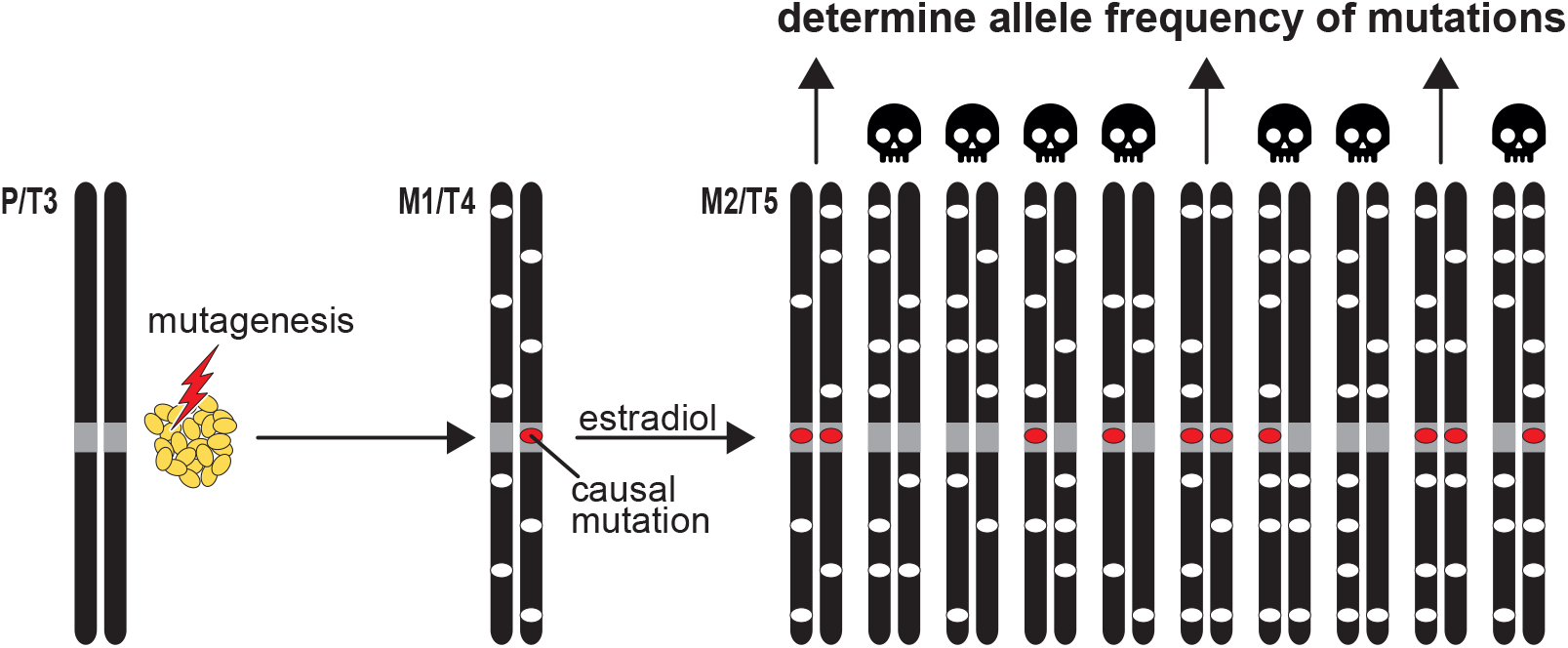
MAD-mapping excludes crosses to expedite isolation of causative mutation. MAD-mapping identifies causal mutations in the M2 generation. The parental transgenic line (P/T3) contains an inducible transgene (not indicated) that triggers systemic cell death (black skull) upon application of estradiol. Seeds of the parental line were mutagenized by EMS treatment (red arrow) producing M1/T4 plants with EMS-induced mutations (ovals). The causal mutation (red oval), that inhibits activity of the inducible transgene is heterozygous in the M1. EMS mutations of a given M1 segregate in M2/T5 descendants. M2 plants that are homozygous for the causal mutation will survive in presence of the inducer chemical. Survivors of a given M2 family are used to generate DNA pools in which the frequency of EMS-induced mutations is determined by next generation sequencing (NGS). The causal mutation will be homozygous in all survivors and thus will be present at a frequency of 1 (100%) in the pool DNA.

### Three *ace* families have distinct mutations in the Arabidopsis *XRN4*/*EIN5* gene

Whole-genome sequencing was performed on the parental line that was originally used for EMS mutagenesis (*Estr:Bs4C-FLAG-GFP*) and three distinct DNA pools composed of survivors from *ace1*, *ace2*, and *ace3* families, respectively. The DNA pools of *ace1*, *ace2*, and *ace3,* contained 38, 18, and 25 M2 survivors, respectively. Paired end sequencing was used with a minimum depth of 150X coverage to determine EMS mutations and their allele frequencies in *ace1*, *ace2*, and *ace3* M2 families (Fig. 4, Fig. S2-S4, Table S1). We detected 60, 150, and 487 EMS mutations specific for the *ace1*, *ace2,* and *ace3* M2 families, respectively. Scanning the pooled genomes of M2 survivors for a selection-induced increase in the frequency of mutant alleles, we identified genomic regions with increased (and locally fixed) mutant allele frequencies on chromosome 1 for all three *ace* mutants (Fig 4). In order to select candidate mutations in these genomic regions that possibly cause the observed cell death suppression phenotype, we limited our search for causal mutations by considering only EMS mutations with an allele frequency of 0.95 or higher. Moreover, our search was restricted to base pair changes that are characteristic to EMS-induced mutations (C to T or G to A). We disregarded mutations that occurred in either the non-coding regions (intronic or untranslated regions) or caused synonymous mutations in the coding regions, thereby focusing on missense and nonsense mutations in coding regions. This narrowed our search down to one candidate gene in *ace1* and *ace2* pools (*AT1G54490.1*), and to three candidate genes in the *ace3* pool (*AT1G54490.1*, *AT1G52940.1*, and *AT1G55110.1*). *AT1G54490.1*, which encodes the EXORIBONUCLEASE 4 (XRN4) / ETHYLENE INSENSITIVE 5 (EIN5) protein (Fig. 4) (20–22), was found to be mutated across all three *ace* families, indicating the functional impact of mutations in this specific CDS on the common phenotype. As one would expect, each M2 family had distinct mutations in the coding region of *AT1G54490.1* (Fig. 5, Fig. S5). The *ace1* mutation lies within the sixth exon, and changes the wildtype aspartate to an asparagine (D to N). The *ace2* mutation is in the eighth exon, and changes the wildtype tryptophan to a premature stop codon (W to *). The *ace3* mutation is found in the third exon of *AT1G54490.1*, and alters the parental or wildtype amino acid of an alanine to a valine (A to V) (Fig. 5).

**Figure 4.**
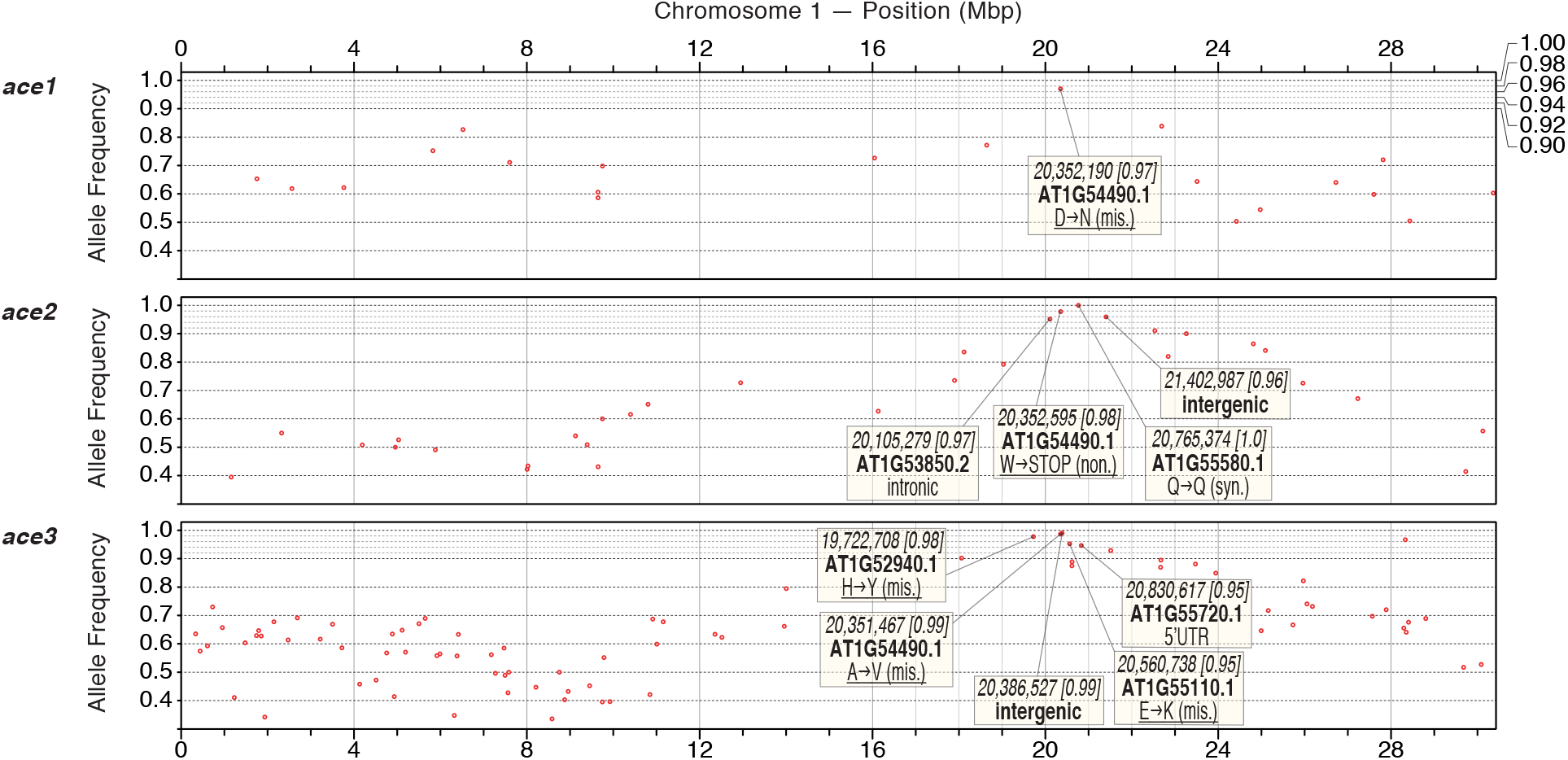
M2 allele frequency-distribution (MAD) mapping identifies mutations in *AT1G54490.1* that suppress Bs4C-dependent cell death. The frequency of EMS-induced mutations (red dots) on chromosome 1 is displayed for *ace1*, *ace2* and *ace3* DNA pools. Boxes provide information on mutations that occur at frequencies of 0.95 or higher. Italic font indicates the chromosomal location of the mutation with the frequency provided in square brackets. Boldface font provides the gene designation. If the mutation is within a gene, the third row reveals the consequences of this mutation. Underlining indicates that the given mutation has likely functional consequences.

**Figure 5.**
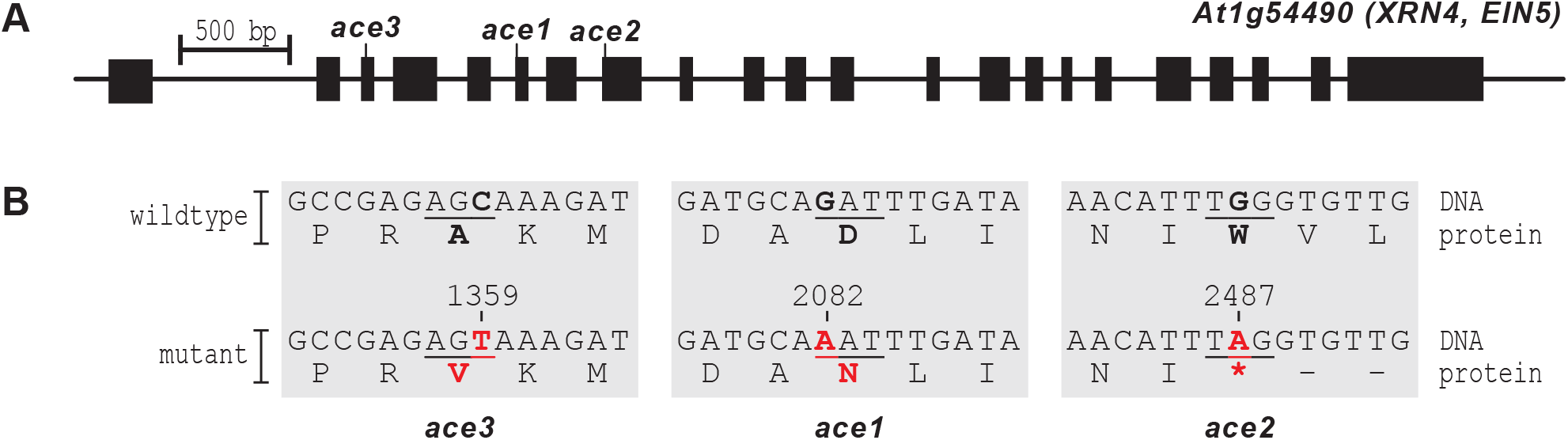
*ace* families harbor distinct mutations in *AT1G54490.1*. **A** | Location of mutations in *ace1*, *ace2*, and *ace3* mutants. Black boxes represent *AT1G54490.1* exons. The location of each causal mutations in *ace3*, *ace1*, and *ace2*, is indicated. Black bar indicates length of 500 bp. **B** | Mutations in *ace* mutants and its consequences at the protein level. Underlined letters indicate the affected codons with the encoded amino acid shown below. Letters in black bold display the base pair or amino acid found in the parental line. Red bold font indicates EMS-induced mutations with the encoded amino indicated below. Numbers above mutated base pairs indicate positions of the mutations within the transcript sequence. An asterisk (*) indicates a translational stop codon.

Altogether, our data suggests that the identified *XRN4/EIN5* mutant alleles abolish Bs4C dependent cell death in Arabidopsis. Moreover, our data demonstrates that MAD-mapping is a highly-efficient approach for identification of causal mutations in M2 mutant families.

## DISCUSSION

We demonstrated that the estradiol-inducible expression of the pepper executor-type R protein Bs4C triggers systemic cell death in transgenic Arabidopsis plants. This conditionally lethal phenotype was used in a forward genetic screen to identify three distinct Arabidopsis *ace* mutants that do not execute Bs4C-dependent plant cell death. We determined the frequency of EMS-mutations in three distinct *ace* M2 families, a process that we designated as MAD-mapping. This identified mutations for all three M2 families within *AT1G54490.1*, which encodes the exoribonuclease XRN4 (22).

As of yet, *XRN4* is the first known genetic component required for cell death triggered by executor-type R proteins. XRN4 is the plant cytoplasmic homolog of yeast and metazoan XRN1, and catalyses degradation of uncapped mRNAs from the 5′ end (23, 24). In a simplistic model, XRN4 could degrade a transcript encoding a negative regulator of the Bs4C-dependent cell death. Absence of functional XRN4 in *ace* mutant plants would presumably cause increased expression of the putative negative regulator and inhibit Bs4C-triggered cell death, being consistent with the observed mutant phenotype.

Recent studies uncovered that PAMP-induced activation of PRRs results in phosphorylation of the DECAPPING 1 (DCP1) protein that in turn, interacts with and activates XRN4 (25). It is assumed that activated XRN4 degrades transcripts encoding positive and negative regulators of PRR-triggered immune reactions. It is therefore conceivable that XRN4 could be a shared regulator of PRR- and executor R protein-triggered immune pathways. Future studies will have to clarify the exact role of XRN4 in Bs4C-dependent cell death reactions, and whether or not XRN4 is also involved in other plant defence pathways.

Forward genetic screens and subsequent isolation of causative mutations by positional cloning is an essential gene discovery tool for elucidation of any kind of biological process in plants (26). The advent of next-generation sequencing technology introduced several innovations into the process of mutation identification, including simultaneous mapping and identification of causal mutations as well as the utilization of isogenic mapping populations (3, 4). Unless allelic groups are available, mutation identification still relies on the time-consuming process of generating numerous segregating populations. Therefore, the workload and time that is needed to establish segregating populations remains a major limitation in forward genetics. We postulated and experimentally validated that the segregation of causal mutations in M2 families, which is regularly used for the initial identification of mutant phenotypes, can already be used to identify causal mutations, ultimately removing the need for tedious generation of segregating populations. Therefore, upon mutagenesis of seeds, only two generations are needed to identify causal mutations. Given the generation time of approximately 8 weeks in Arabidopsis, it essentially takes less than one year to identify causal mutations via MAD-mapping. While we demonstrated the feasibility of MAD-mapping in the model plant Arabidopsis, the concept could be applicable to any plant and even non-plant species.

We combined MAD-mapping with a conditionally lethal screen (Fig. 3); and a benefit of this combination is that it can be carried out at the seedling stage. Accordingly, large numbers of mutants can be studied in a short time and the need for space remains quite limited. Plant defence reactions typically rely on the execution of cell death reactions, and as a result, several conditionally lethal screens have been conducted in the past to study plant R proteins and to identify signalling components of R pathways (27–29). These screens depend on inducible promoters that typically contain constitutively expressed elements. For example, the estradiol-inducible system that we used contains the constitutively expressed synthetic transcription factor XVE that is activated by estradiol (Fig. 1). It has been noted in the past, that estradiol-inducible transgenes lose inducibility throughout generations (30). This phenomenon typically starts in the T4 and T5 generations and is likely the consequence of transgene silencing. We identified causal mutations in the M2 generation, which corresponds to the T5 generation (Fig. 3). In previous studies, we used the estradiol-inducible system in a conventional forward genetic screen and established conventional F2 mapping populations, which corresponds to the T7 generation, to identify causal mutations for given M2 survivors. However, we did not observe the expected segregation of cell death in F2 individuals and ultimately could not identify causal mutations by this approach, possibly caused by transgene silencing in the F2/T7 mapping generation. In MAD-mapping, phenotypic identification and isolation are both carried out in the M2 generation, essentially overcoming the problem of gene silencing that possibly occurs in mapping populations derived from a single transgenic M2 plant.

While the principle of MAD-mapping is broadly applicable, it cannot be carried out on bulked M2 populations since it is based on the analysis of individual M2 families. Accordingly, after EMS mutagenesis, each M1 plant must be harvested individually to generate a collection of M2 families. Similarly, each M2 family must be studied individually for phenotypic changes. Although MAD-mapping is generally time-saving, it is more laborious in the harvesting and screening phase than conventional screens that are typically based on bulked M2 seeds. On the upside, however, screening of separate M2 families offers the possibility for recovering mutations that are infertile when homozygous via the heterozygous siblings of the mutant plants. Moreover, this strategy guarantees the independence of mutants isolated from distinct M2 families. In the long run, while the analysis of M2 families is more laborious than analysis of bulked M2 seeds, the benefits of MAD mapping vastly overcome the short-term extra work that is required.

Overall, we envision that the ease and speed of MAD-mapping will substantially increase the attraction of forward genetic approaches and it stands to reason that MAD-mapping will make a major contribution towards the elucidation of biological pathways in the near future.

## MATERIALS AND METHODS

### Plant material and growth conditions

*Arabidopsis thaliana* plant material used in this study: Col-0, *Estr:Bs4C-FLAG-GFP, Estr:GFP-GUS, ace1 family, ace2 family, ace 3 family.* For the seedling growth assay, seeds were sterilized using 80% ethanol and 0.05% Triton X-100 solution, and left to stratify in the darkness at 4 C for two days on ½ MS plates (0.43% (w/v) MS Salts (Gibco), 1% (w/v) Sucrose, 0.05% MES, pH 5.8) containing 200 μg/mL Cefotaxim. Seeds were put to long day (16hr light/8hr dark) at 20 °C in light and 18 °C in dark for four days. On the fourth day, seedlings were transplanted to 48-well plate, each well containing either 20 μM estradiol, or mock treatment (1% (v/v) DMSO), and left for 10 more days. On the 14^th^ day, seedling growth was analysed. For seedling immunoblot detection, four 14 day old seedlings of each genotype were placed in either 20 μM estradiol or 1% DMSO, vacuum infiltrated, and left at room temperature for 24 hours. Samples were then flash frozen and used for immunodetection, as described below. For EMS mutagenesis, approximately 200 mg of *Estr:Bs4C-FLAG-GFP* Arabidopsis seeds were allowed to swell in water for 3 days. Afterwards, these seeds were incubated in 50 mL of 0.3% EMS solution for 6 hours, shaking. The seeds were then transferred to a Nalgene Filter Unit and washed six times with water. The seeds were then resuspended in 0.1% phytoagar and sowed on soil. After 10 days, the seedlings were transplanted into individual pots in an outdoor greenhouse (16hr light/8hr dark, temperature minimum 18° C, no humidity control). These individual plants now generated the M1 population. After 6 weeks, seeds from each individual M1 plant were harvested, creating 4000 individual M2 families. For the screening of *ace* families, 100 seeds of each of the 4000 M2 families were placed in a 96 well plate, and were gas sterilized (80 mL NaClO and 3 mL 32% HCl solution) overnight. The following day, 200 μL of 1% phytoagar was placed in each well, sealed, and left to stratify at 4 °C in the dark for 2 days. The seeds were then plated on ½ MS plates (described previously) containing 20 μM estradiol, and put to long day. Plates were left for 14 days, suppressing families were selected, and survivors were transplanted to soil. Five week old plants were used for immunodetection by taking three 4 mm punches and vacuum infiltrating them with 20 μM estradiol, and letting them sit for 24 hours before flash freezing and going forward with immunodetection (as described below). To sequence the transgene, gDNA was collected from leaf tissue, PCR amplified, and sent for Sanger sequencing.

### Plasmid construction

For *Estr:Bs4C-FLAG-GFP* and *Estr:GFP-GUS* T-DNA constructs coding sequences of *Bs4C*, *3xFlag*, *GFP* and *uidA* were PCR amplified and cloned via GoldenGate cloning into pENTR CACC-AAGG. Resulting pENTR-Bs4C-FLAG-GFP and pENTR-GFP-GUS were used in LR reaction together with pER10-GW generating pER10-Bs4C-FLAG-GFP and pER10-GFP-GUS.

### Transgenic lines

*Estr:Bs4C-FLAG-GFP and Estr:GFP-GUS* were generated using *Agrobacterium* GV3101 containing pER10-Bs4C-FLAG-GFP or pER10-GFP-GUS in a floral dip method. Transgenic *A. thaliana* were selected with Kanamycin on ½ MS plates.

### Genomic DNA extraction

Approximately 100-150 mg of leaf material was collected. 600 μL of CTAB buffer (100 mM Tris-HCl pH 8.0, 20 mM EDTA, 1.4 M NaCl, 2% (w/v) cetyltrimethyl ammonium bromide) was added, and homogenized using a vortex. Samples were incubated at 65 °C for 30 minutes. Heated samples were spun down at room temperature. 500 μL of the supernatant was transferred to a new tube. 2.5 μL of RNAse A (10 mg/mL, ThermoFisher) was added, and gently vortexed, and incubated at 37 °C for 30 min. 500 μL of chloroform was added, and mixed. Samples were spun down at room temperature, and 450 μL of the aqueous phase was added to a new tube. 450 μL of 100% isopropanol was added, and gently mixed. The tubes were then spun down until a pellet formed, and the supernatant was discarded. 500 μL of 70% ethanol was added, mixed, spun down at room temperature, and then the supernatant was discarded. This was repeated twice. The pellet was then dried at 35 °C. The dried pellet was dissolved in 35 μL of 10 mM Tris-HCl pH 8.0, and quantified using a Qubit (ThermoFisher). These samples were then sent for NGS.

### Next generation sequencing and mapping populations

Raw reads of each sample were aligned to Col-0 reference genome (The Arabidopsis Genome Initiative 2000; www.arabidopsis.org) using *GenomeMapper* (31), after which short-read alignments were corrected for read-pair information and consensus bases were called with *shore* (31). After removing common SNPs between each mutant and the parental line, the causal mutation of each mutant was predicted by analysing allele frequencies with SHOREmap v3.0 (2, 32).

### Immunoblotting

Samples were flash frozen and then ground to a fine powder. 50 μL of SDS loading buffer (50 mM Tris-HCL pH 6.8, 100 mM DTT, 2% SDS, 0.1% bromophenol blue, 10% glycerol) was added, and boiled at 95 C for 10 min. Samples were loaded onto a SDS-polyacrylamide gel (4% stacking, 10% resolving), and then transferred to a PVDF-membrane (BioRad). Samples were blocked in 5% milk/1X TBST (50 mM Tris-HCl, 150 mM NaCl, 0.05% Tween-20), and anti-bodies were then applied. Anti-FLAG primary antibody (F1804, Sigma-Aldrich) raised in mouse, at 1:5000 dilution, was used, shaking overnight. The next day, membranes were washed with 1X TBST (50 mM Tris base, 150 mM NaCl, 0.05% (v/v) Tween-20), and the anti-mouse-HRP secondary antibody (A9044, Sigma-Aldrich) was used at a 1:2500 dilution, and incubated for 2 hours. Anti-GFP-HRP conjugated primary antibody (SC-9996, SantaCruz) at 1:2500 dilution was used, and incubated for 2 hours. The blot was washed 3 times with 1X TBST, and once with 1X TBS (1X TBST, but no Tween-20 added). The Clarity ECL Substrate (BioRad) and the Amersham™ Imager 600 (GE Life Sciences) machine were used for imaging. All membranes were stained with Ponceau.

## ACKNOWLEDGEMENTS

The work was supported by the Deutsche Forschungsgemeinschaft (DFG) [SFB 1101 to T. Lahaye (project D08) and F. El Kasmi (project D09) and LA 1338/7-1 to T. Lahaye]. Research by K. Schneeberger was funded by the DFG under Germany’s Excellence Strategy - EXC 2048/1 - 390686111, and the European Research Council (ERC) Grant “INTERACT” (802629). We thank E. S. Ritchie and A. Strauß for helpful comments on this manuscript, and S. Üstün for insightful discussions. We would like to thank A. Dressel, N. Gallas, P. Gouguet, P. Lutz, T. Phan, E. S. Ritchie, K. Schenstnyi, L. Schmaltz, S. Schade, A. Strauß, D. Wu, and Y. You for their help with separating and collecting M1 plants, as well the ZMBP gardeners for taking care of these plants.

## SUPPLEMENTAL FIGURES

**Figure S1.**
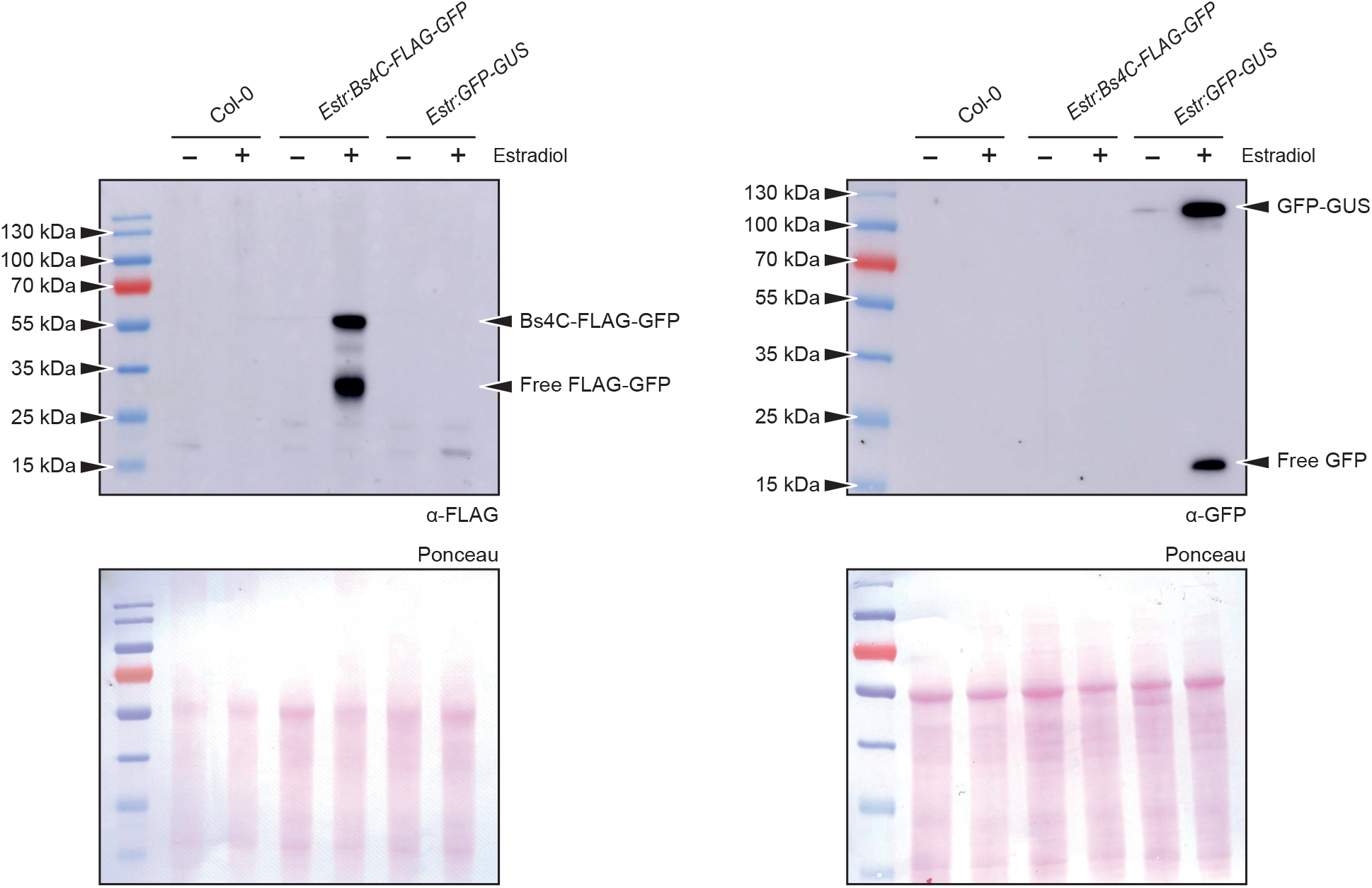
Transgenic lines containing inducible transgenes express transgene-encoded proteins in an estradiol-dependent fashion. Immunoblot analysis using anti-FLAG and anti-GFP antibody of tissue from two week old Arabidopsis seedlings of depicted genotypes (Col-0, *Estr:Bs4C-FLAG-GFP*, *Estr:GFP-GUS*). Plants were incubated for 24 hours in liquid media either containing estradiol or lacking estradiol (mock). Ponceau stained membrane serves as a loading control. Molecular mass markers are indicated by triangles.

**Figure S2.**
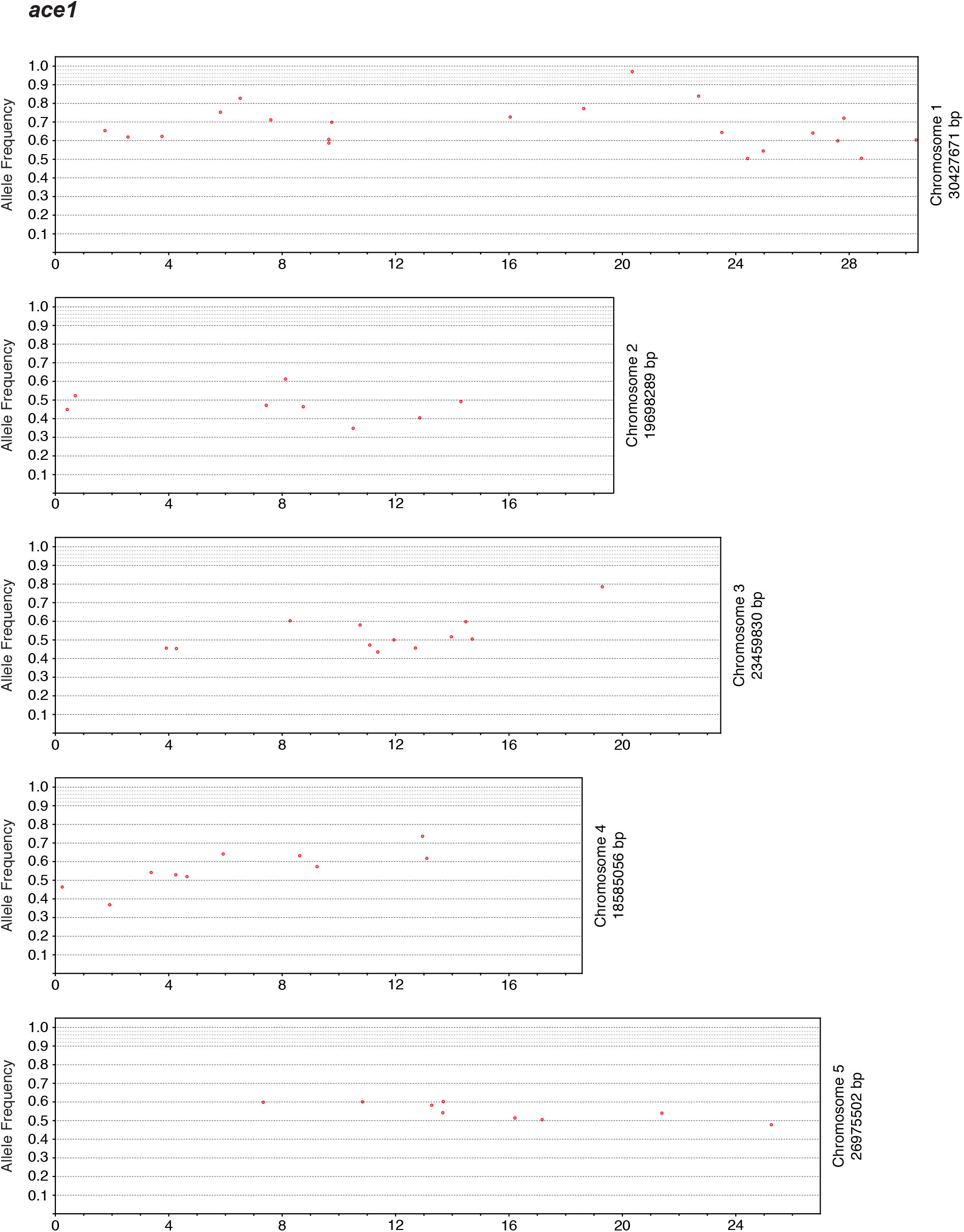
M2 allele frequency-distribution in the *ace1* mutant family. The frequency and position of EMS-induced mutations (red dots) on chromosome 1-5 are displayed.

**Figure S3.**
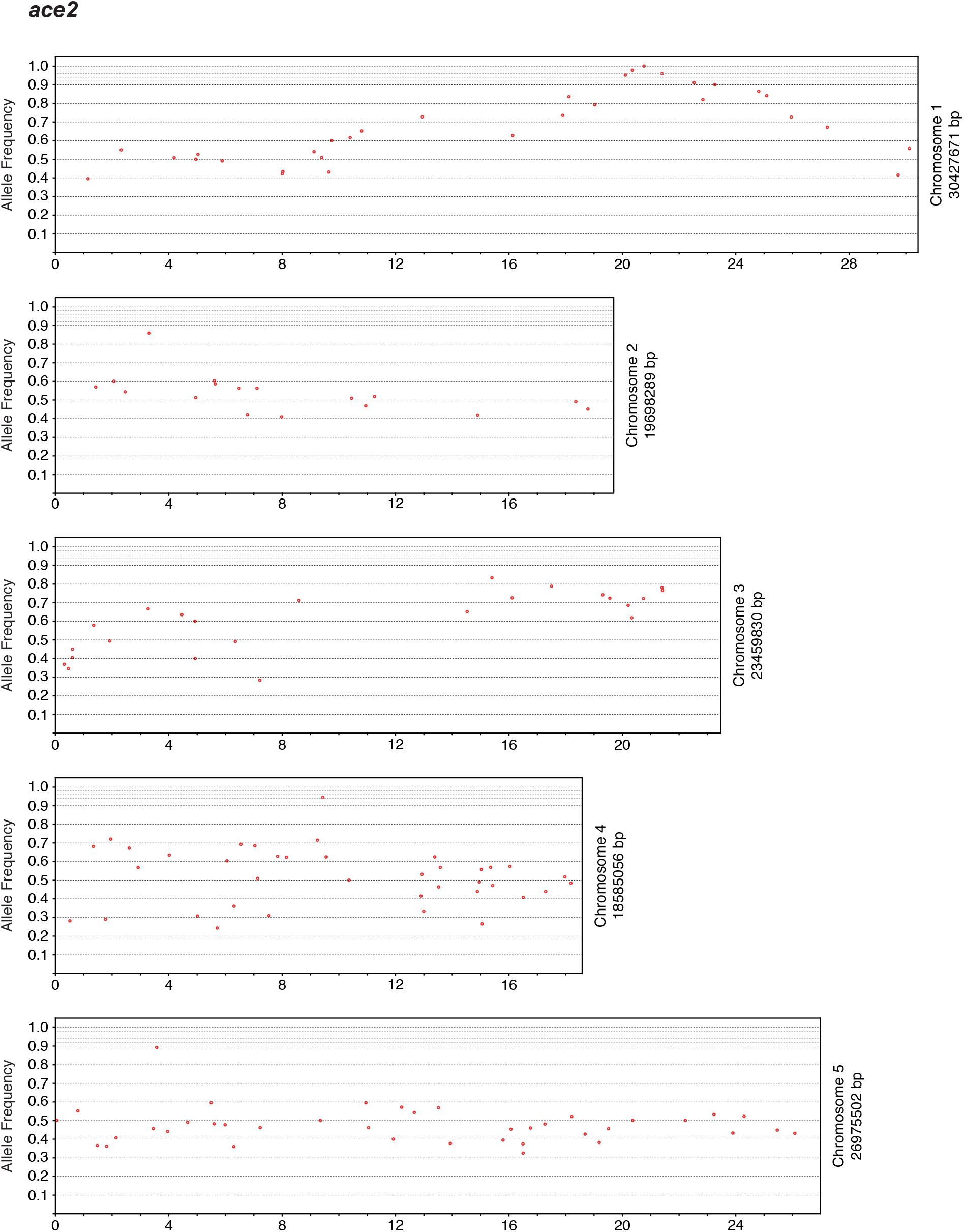
M2 allele frequency-distribution in the *ace2* mutant family. The frequency and position of EMS-induced mutations (red dots) on chromosome 1-5 are displayed.

**Figure S4.**
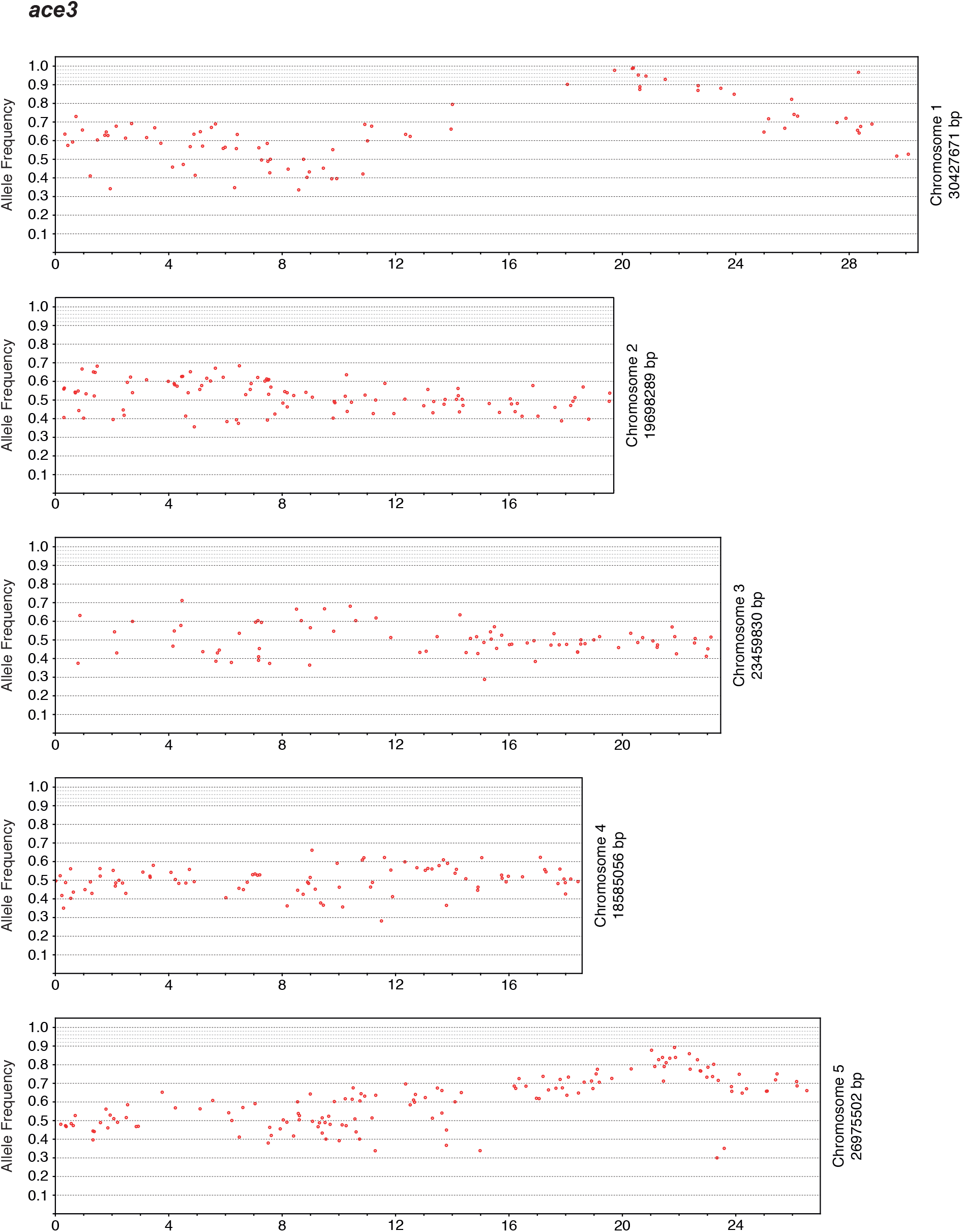
M2 allele frequency-distribution in the *ace3* mutant family. The frequency and position of EMS-induced mutations (red dots) on chromosome 1-5 are displayed.

**Figure S5.**
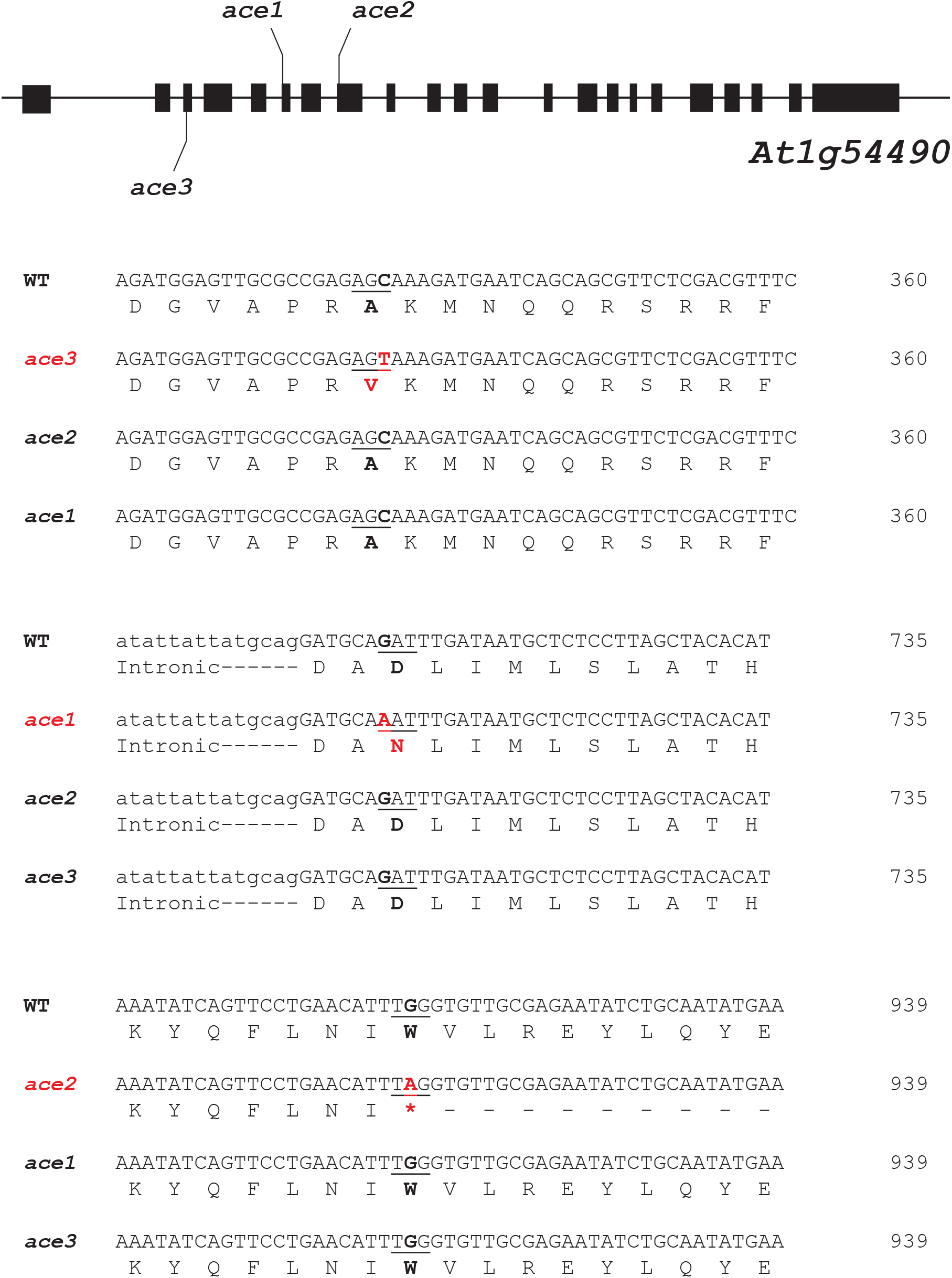
Mutations in *ace* families are independent from one another. Underlined letters indicate the 3 letter codon corresponding to the amino acid directly below. Letters in black bold display reference base pair or amino acid found in the parental line, and letters in red bold designate altered base pair or amino acid found in indicated mutant family.

## SUPPLEMENTAL TABLES

Table S1 – EMS-induced mutations with increased allele frequency in Arabidopsis *ace* mutants.

**Table S1.** Allele frequencies of EMS-induced SNPs in three *ace* families.

| Family | Position | Ref./Alt. | Cover. | Allele freq. | Gene ID | Feature | CDS position | Effect | Change |
| --- | --- | --- | --- | --- | --- | --- | --- | --- | --- |
| *ace1* | 18,641,436 | G/A | 81 | 0.77 | — | intergenic |  |  |  |
|  | 20,352,190 | G/A | 98 | 0.97 | <u>AT1G54490.1</u> | CDS | 706 | Nonsyn | D/N |
| *ace2* | 18,117,826 | G/A | 61 | 0.85 | AT1G48970.1 | 5' UTR |  |  |  |
|  | 19,031,508 | G/A | 61 | 0.87 | — | intergenic |  |  |  |
|  | 20,105,279 | G/A | 59 | 0.97 | AT1G53850.1 | intronic |  |  |  |
|  | 20,105,279 | G/A | 59 | 0.97 | AT1G53850.2 | intronic |  |  |  |
|  | 20,352,595 | G/A | 44 | 0.98 | <u>AT1G54490.1</u> | CDS | 911 | Nonsyn | W/* |
|  | 20,765,374 | G/A | 59 | 1 | AT1G55580.1 | CDS | 1269 | Syn | Q/Q |
|  | 21,402,987 | G/A | 47 | 0.96 | — | intergenic |  |  |  |
| *ace3* | 18,062,237 | C/T | 101 | 0.9 | AT1G48840.1 | intronic |  |  |  |
|  | 19,722,708 | C/T | 129 | 0.98 | AT1G52940.1 | CDS | 478 | Nonsyn | H/Y |
|  | 20,351,467 | C/T | 146 | 0.99 | <u>AT1G54490.1</u> | CDS | 329 | Nonsyn | A/V |
|  | 20,386,527 | C/T | 102 | 0.99 | — | intergenic |  |  |  |
|  | 20,560,738 | C/T | 139 | 0.95 | AT1G55110.1 | CDS | 1036 | Nonsyn | E/K |
|  | 20,613,630 | C/T | 98 | 0.88 | — | intergenic |  |  |  |
|  | 20,616,883 | C/T | 96 | 0.89 | — | intergenic |  |  |  |
|  | 20,830,617 | C/T | 123 | 0.95 | AT1G55720.1 | 5' UTR |  |  |  |
|  | 21,512,799 | C/T | 130 | 0.93 | AT1G58100.1 | CDS | 1087 | Nonsyn | G/S |
|  | 21,512,799 | C/T | 130 | 0.93 | AT1G58100.2 | CDS | 1015 | Nonsyn | G/S |

SNPs identified on Arabidopsis chromosome 1 in *ace1*, *ace2* and *ace3* M2 families, stating the position, the reference base (Ref.), identified altered base (Alt.), the coverage (Cover), allele frequency (Allele freq.), the annotated Gene ID, the feature within the gene, the coding sequence position (CDS position), the effect of the mutation, and the resulting amino acid change (Change). SNPs that were below an allele frequency of 0.77 are not listed. Underlined font indicates the gene that was identified in all three *ace* families.

